# Visual Perceptual Learning of Feature Conjunctions Leverages Non-linear Mixed Selectivity

**DOI:** 10.1101/2022.10.04.510801

**Authors:** Behnam Karami, Caspar M. Schwiedrzik

## Abstract

Visual objects are often defined by multiple features. Therefore, learning novel objects entails learning conjunctions. Visual cortex is organized into separate compartments, each of which is devoted to processing a single feature. A prime example of this is are neurons purely selective to color and orientation, respectively. However, neurons that jointly encode multiple features (mixed selectivity) also exist across the brain and play critical roles in a multitude of tasks. Here, we sought to uncover the optimal policy that our brain adapts to achieve conjunction learning using these available resources. 59 human subjects practiced orientation-color conjunction learning in four psychophysical experiments designed to nudge the visual system towards using one or the other resource. We find that conjunction learning is possible by linear mixing of pure color and orientation information, but that more and faster learning takes place when pure and mixed selectivity neurons are involved. We also find that learning with mixed selectivity confers advantages in performing an untrained “exclusive or” (XOR) task several months after learning the original conjunction task. This study sheds light on possible mechanisms underlying conjunction learning and highlights the importance of learning by mixed selectivity in such accounts.

## Introduction

Visual objects are often defined by conjunctions of multiple features. This means that learning to discriminate visual objects requires faithful learning of these conjunctions. For example, to become a bird expert who successfully differentiates among subspecies of finches, it is necessary to be highly sensitive to the color and angle of the beak together. To date, studies in visual perceptual learning have almost exclusively focused on single feature learning and therefore, a plausible framework to explain conjunction learning is still lacking.

Decades of research in neurophysiology of the visual system have established that cortical representations of visual features diverge from very early stages of visual processing onwards. Particularly well known is the divergence of color and orientation, which are represented in separate anatomical compartments and processed in parallel processing streams (1–11). However, more recent work has pointed out that this segregation is not absolute, but that neurons which code jointly for orientation and color exist along the entire visual ventral stream (albeit, perhaps, in low numbers) (12–17). Taken together, these studies point to the existence of three groups of neurons: purely orientation selective, purely color selective, and neurons with mixed selectivity between color and orientation.

How can conjunction learning be achieved by means of these neuronal resources (Fig. 1a)? One strategy, and perhaps the simplest one, is to improve processing of each constituent feature of a conjunction separately, and to linearly combine them at the output level. This strategy is completely sufficient for linearly separable tasks and makes optimal use of pure selectivity neurons (which are seemingly larger in number), while disregarding mixed selectivity neurons. Another plausible strategy would be to rely only on mixed selectivity neurons. Although fewer in number, these neurons have the advantage that they explicitly convey task-relevant information on the conjunction level. Furthermore, recent research has pointed out computational advantages of using (nonlinear) mixed selectivity representations in a variety of tasks (18). Under these two strategies only a set of highly sensitive neurons that provides the most relevant information for the task is read out, commonly referred to as “precision pooling” (Fig. 1a, top panels) (19,20). An alternative, third strategy is “global pooling” which posits that all active sensory neurons contribute to the decision regardless of their sensitivity (21–23). It is only within this framework that both pure and mixed selectivity neurons would be jointly engaged in learning (Fig. 1a, bottom panel).

**Fig. 1.**
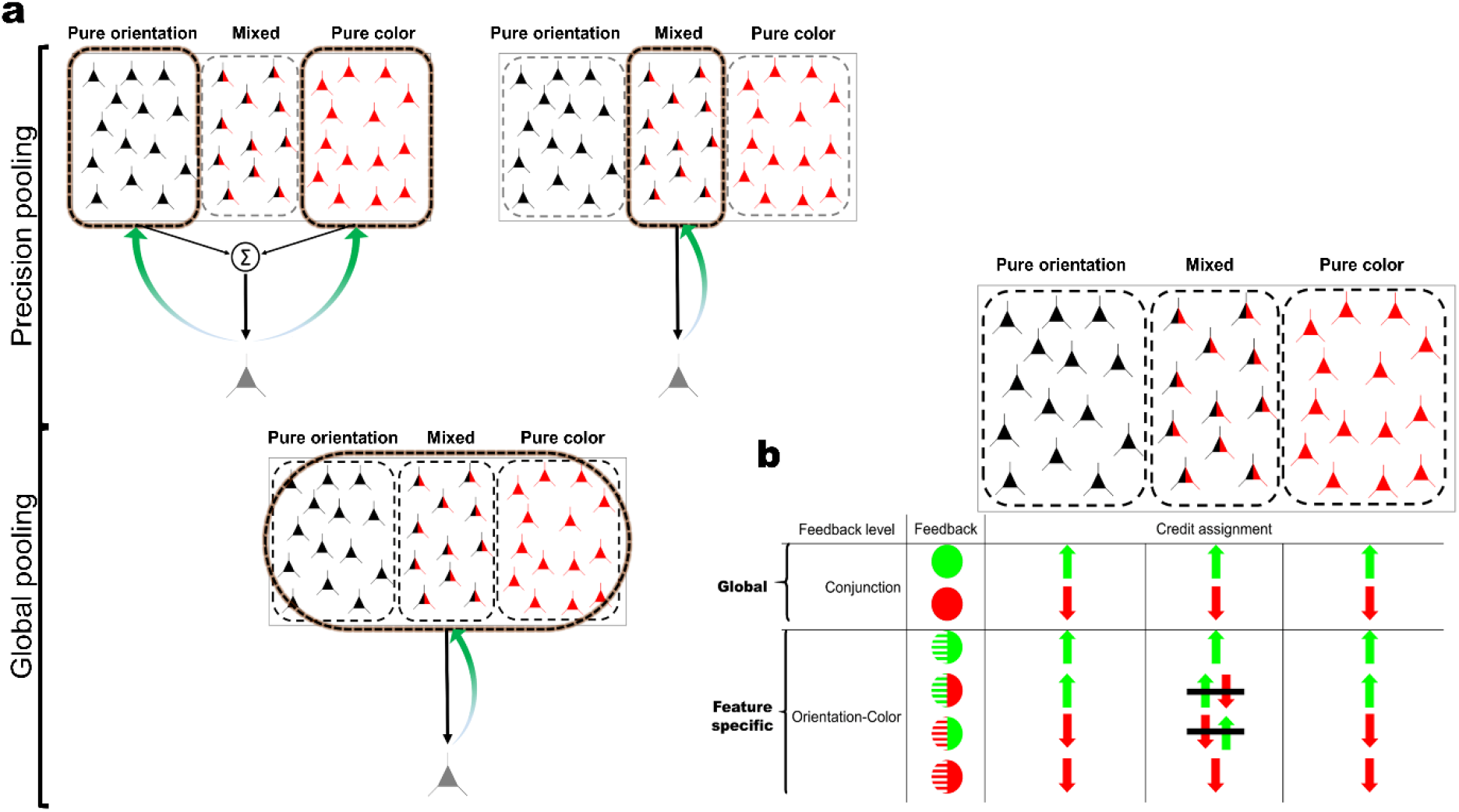
Precision vs. global pooling and credit assignment. **a,** Precision pooling: in precision pooling, only a subset of neurons of a population is read out. For conjunction learning, one option is to read out only neurons purely tuned to color and to orientation, respectively. Their output can be summed to give rise to conjunction information (top left panel). Alternatively, only neurons with mixed selectivity for color and orientation can be read out. These neurons provide information about conjunctions directly and explicitly (top-right panel). Global pooling: In global pooling, all neurons are read out regardless of their tuning properties. Hence, neurons purely tuned to color and to orientation, as well as neurons with mixed selectivity for both features, contribute to conjunction learning. **b,** Credit assignment regimes in conjunction learning with global or feature-specific feedback. When feedback is global (on the conjunction level), the three relevant resources, pure orientation, pure color, and mixed selectivity, are up-/down-weighted together. However, when feature-specific feedback is provided, credit is only consistently assigned to pure selectivity neurons. When one feature is correct and the other incorrect, credit cannot be unambiguously assigned to neurons with mixed selectivity (signified by black lines on up and down-weighting arrows in the middle column). Hence, their weights are updated less frequently (or consistently) than those of pure selectivity neurons, i.e., only when the response is correct or incorrect on both features.

To appraise these strategies, we conducted a set of experiments in which participants were trained for several days to discriminate orientation-color conjunctions. In our main experiments, we used a linearly separable task, where all the above strategies could be reasonably employed. To nudge the visual system towards one or the other strategy, we experimentally manipulated the feedback that subjects received (Fig. 1b): this was either informative on the conjunction level, i.e., subjects had to be correct in both dimensions (global feedback); or on the feature level, i.e., subjects received separate feedback for either dimension (feature-specific feedback). We reasoned that global feedback favors the incorporation of mixed selectivity neurons, while feature-specific feedback favors pure selectivity because only global feedback leads to consistent weight updates of mixed selectivity neurons.

We found that conjunction learning is possible under both feedback conditions. Intriguingly, however, global feedback resulted in more and faster learning than feature-specific feedback both on single-feature and conjunction levels. This was accompanied by non-linear integration of color and orientation information and better transfer to a non-linear exclusive or (XOR) task in the global feedback condition. Control experiments showed that the relative disadvantage of learning with feature-specific feedback was not due to higher cognitive load or noisy credit assignment. Taken together, this indicates that global feedback favors the integration of mixed selectivity neurons, enabling the computational advantages that come with this form of tuning. However, an additional transfer task in which we presented the stimuli at a new location after training also revealed a contribution of pure selectivity neurons even in the global feedback condition, most consistent with global pooling.

Overall, our findings suggest that when learning conjunctions, the brain makes use of all its computational resources instead of selectively reading out specific neurons. This capacity can be unlocked by providing global feedback, which may seem less informative than feature-specific feedback at first sight but favors the incorporation of non-linearly mixed selectivity neurons and unlocks their computational benefits even when this is not strictly necessary for the task at hand.

## Results

### Conjunction learning (Experiments 1 and 2)

To uncover the optimal strategy to learn feature conjunctions, we designed two experiments in which we trained subjects for four days on an orientation-color conjunction discrimination task. Color and orientation difficulty levels were individually determined per subject in a pre-measurement (Fig. 2a-d, also see Methods). While the conjunction task was identical in both experiments, we provided different choice feedback to the subjects, building upon the idea that feedback information can modulate population of neurons to be engaged in learning (Fig. 1b). In Experiment 1 (*n*=19), choice feedback was informative on the conjunction level (global feedback), i.e., correct feedback followed a response only if it was correct on both dimensions (color and orientation). Otherwise, error feedback was provided at the end of the trial (irrespective of which feature was incorrect). In Experiment 2 (*n*=18), we provided choice feedback regarding the accuracy on each dimension separately (i.e., whether color or orientation were correct, respectively). Here, when one feature is correct and the other incorrect, credit cannot be unambiguously assigned to neurons with mixed selectivity. We hypothesized that if conjunction learning is a result of linear mixing of pure orientation and color selectivity neurons, it should not be affected by our feedback manipulation. On the other hand, if mixed selectivity neurons (which explicitly encode conjunctions) play a role, learning should suffer from excluding or impeding the contribution of these neurons.

**Fig. 2.**
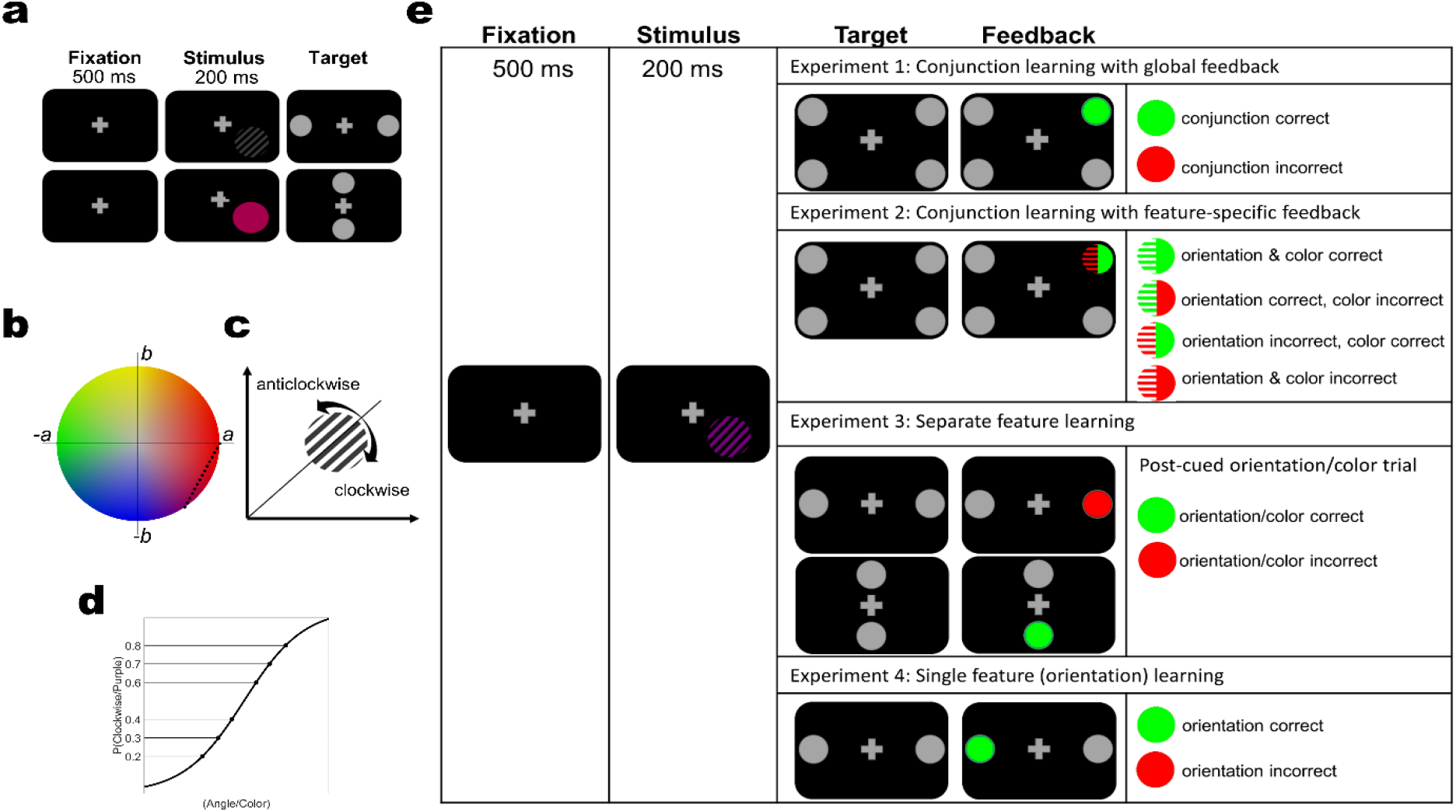
Stimuli and experimental design. **a,** Pre/post training orientation and color discrimination tasks (top and bottom rows, respectively). In separate runs, a 500 ms fixation period was followed by a 200 ms presentation of a grating or colored disc, respectively. Response targets were presented on the left/right or top/bottom of the fixation cross the for orientation and color task, respectively. Subjects responded by directing their gaze to the target. In all experimental groups, these tasks were conducted before training to determine individual difficulty levels, and repeated after training course to establish learning effects on individual features. **b,** Colors were uniformly sampled from the red-purple spectrum of the LAB color space (black dots signify color levels). **c,** Subjects had to determine whether a grating was tilted clockwise or anti-clockwise with reference to the diagonal (45°) in the orientation discrimination tasks. **d,** Six individual difficulty levels for color and orientation discrimination, respectively, were determined from the psychometric curves of the two tasks. **e,** In the main experiments, a 500 ms fixation period was followed by a 200 ms presentation of the stimuli (as in the pre/post measurements). The stimuli were chromatic gratings with individually determined orientation/color difficulty levels. In Experiments 1 and 2, subjects were trained on orientation-color conjunction discrimination. In Experiment 1, we provided global feedback on each trial, i.e., feedback informed about accuracy on the conjunction level (feedback was provided by changing the color of the chosen target; green signifies correct and red incorrect responses). In Experiment 2, we provided feature-specific feedback (e.g., orientation correct, color incorrect). In Experiment 3, subjects were exposed to the same stimuli as in Experiments 1 and 2, but were post-cued to respond only to one feature in a given trial to encourage separate feature learning. The location of the targets (top/bottom or left/right) indicated which feature to respond to. The order of features was pseudo-randomized. Feedback was provided for the given response. In Experiment 4, subjects were trained on a discriminating a single feature, orientation. Color varied from trial to trial but had to be ignored. Color difficulty levels were not individually determined but chosen to maximize between-level distances in the sampled LAB space to ensure high saliency in the color dimension.

To evaluate these hypotheses, we computed the Learning Index (LI), which quantifies learning relative to baseline performance (see Methods). A mixed non-parametric ANOVA showed that participants in both experiments learned to discriminate conjunctions, as evidenced by significant block-by-block improvements in LI (Experiment 1: *F*(6,126)=32.855, *p*<10^-15^, *η^2^*=0.61, Experiment 2: *F*(6,119)=27.397, *p*<10^-15^, *η^2^*=0.58). Interestingly, global feedback quickly led to significantly faster and overall more learning than feature-specific feedback (Fig. 3a), as shown by a significant experiment-by-block interaction (Experiment: *F*(1,35)=3.0793, *p*=0.08, *η^2^*=0.0809, Block: *F*(6,210)=197.4323, *p*<10^-15^, *η^2^*=0.8495, Experiment×Block: *F*(6,210)=4.5752, *p*=0.0002, *η^2^*=0.1156). Comparing LIs per time point relative to baseline between experiments we found that from block 2 onwards, global feedback outperformed feature specific feedback (Experiment 1 vs. Experiment 2, Baseline - block1: *z*=-10.4591, d=-0.2614, *p*=0.5833, block2: *z*=-59.26, *d*=-1.48, *p*=0.0021, block3: *z*=-70.3246, *d*=-1.7578, *p*=0.0003, block4: *z*=-64.383, *d*=-1.6093, *p*=0.0008, block5: *z*=-67.1111, *d*=-1.6775, *p*=0.0005, test: *z*=-52.576, *d*=-1.3142, *p*=0.0062).

**Fig. 3.**
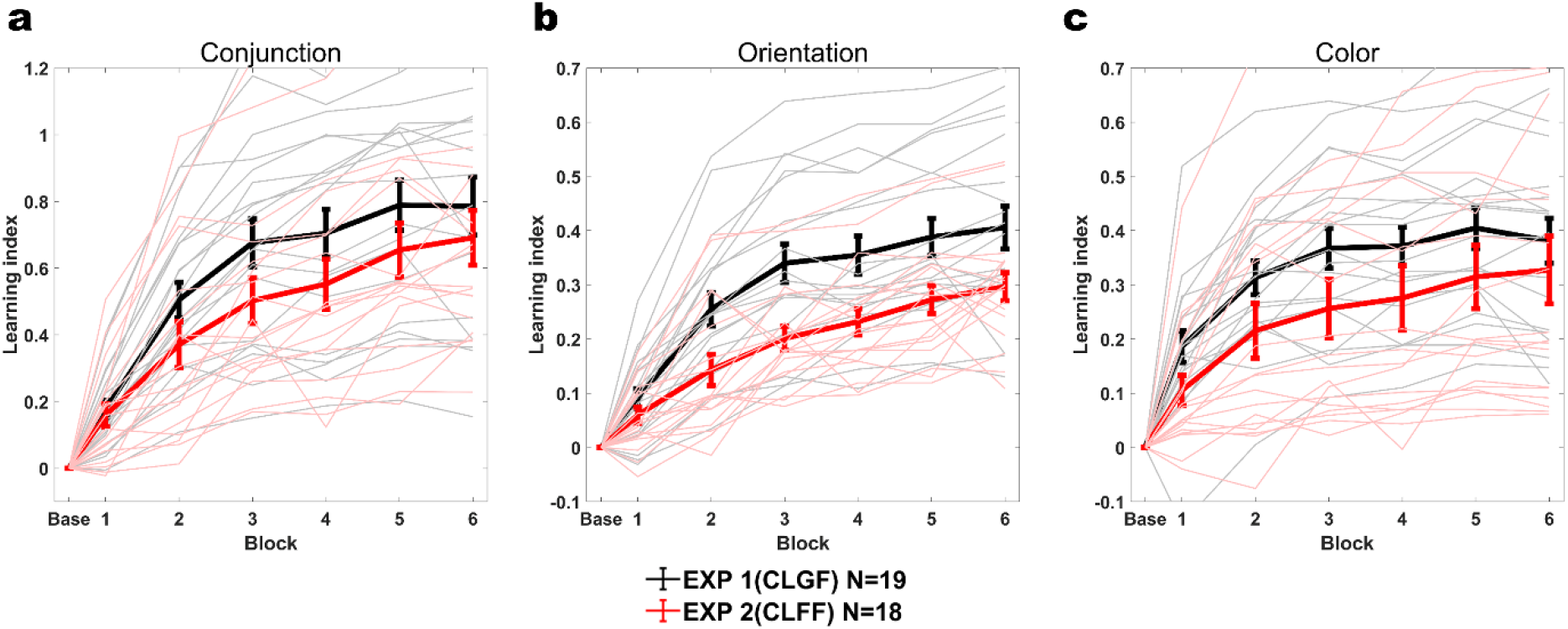
Learning indices (LI) for conjunction learning with global and feature-specific feedback (Experiments 1 and 2). **a,** Conjunction LIs across training blocks. The initial 144 trials of the learning phase were taken as baseline (Base). Thick lines are average LIs across subjects, light thin lines are individual subjects’ LIs. Global feedback (CLGF) in Experiment 1 resulted in more learning than feature-specific feedback (CLFF) in Experiment 2 (Experiment×Block *F*(6,210)=4.5752, *p*=0.0002, *η^2^*=0.1156) **b,** Orientation LIs. These values were computed based on the accuracy in the orientation dimension during conjunction learning. Global feedback (CLGF) in Experiment 1 resulted in more learning than feature-specific feedback (CLFF) in Experiment 2 (Experiment×Block *F*(6,210)=8.7148, *p*<10^-7^, *η^2^*=0.1994) **c,** Color LIs. These values were computed based on the accuracy in the color dimension during conjunction learning. Global feedback (CLGF) in Experiment 1 resulted in more learning than feature-specific feedback (CLFF) in Experiment 2 (Experiment×Block *F*(6,210)=3.9228, *p*=0.001, *η^2^*=0.1008). Error bars represent the standard error of the mean.

We also performed similar comparison on the level of individual features, separately assessing accuracy for color and orientation on the conjunction task (Fig. 3b, c). Here, we found that the advantage of global over feature-specific feedback also held, and was especially pronounced for orientation (Orientation, Experiment: *F*(1,35)=9.7644, *p*=0.0036, *η^2^*=0.2181, Block: *F*(6,210)=178.9749, *p*<10^-15^, *η^2^*=0.8363, Experiment×Block: *F*(6,210)=8.7148, *p*<10^-7^, *η^2^*=0.1994; Color, Experiment: *F*(1,35)=3.8318, *p*=0.0582, *η^2^*=0.0987, Block: *F*(6,210)=114.9568, *p*<10^-15^, *η^2^*=0.7672, Experiment×Block: *F*(6,210)=3.9228, *p*=0.001, *η^2^*=0.1008). Moreover, contrasting LIs between experiments block-by-block yielded similar results as we had observed for conjunctions (Orientation: Experiment 1 vs. Experiment 2, Baseline - block1: *z*= −32.43, *d*=-0.7386, *p*=0.1138, block2: *z*=-59.26, *d*=-1.48, *p*<10^-6^, block3: *z*=-115.25, *d*=-2.6247, *p*<10^-7^, block4: *z*=-96.23, *d*=-2.1916, *p*<10^-5^, block5: *z*=-93.14, *d=-* 2.1212, *p*<10^-5^, test: *z*=-85.39, *d*=-1.9448, *p*<10^-4^; Color: Baseline - block1: *z*=-53.32, *d*=-1.3414, *p*=0.0043, block2: *z*=-67.02, *d*=-1.686, *p*=0.0004, block3: *z*=-76.12, *d*=-1.9149, *p*=0.0001, block4: *z*=-72.13, *d*=-1.8145, *p*=0.0001, block5: *z*=-65.64, *d*=-1.6513, *p*=0.0005, test: *z*=-54.36, *d*=-1.3675, *p*=0.0037). Because LI is computed relative to baseline performance, these results cannot be due to baseline difference between the two learning groups.

If conjunction learning relied on purely color and orientation selective neurons, respectively, one may intuitively assume that feature-specific feedback should lead to better or at least equal performance as global feedback, especially on the level of individual features. Yet, our results point to the opposite conclusion. What could be behind this seemingly counterintuitive finding? We hypothesized that an involvement of neurons with non-linear mixed selectivity between features might account for our results. Non-linear mixed selectivity is thought to increase coding reliability and efficiency not only on the level of conjunctions but also on the level of constituent features (18).

To probe if and to what extent non-linear mixing accounted for the observed effect, we quantified the degree of non-linearity in the combination of color and orientation in our data. To this end, we applied a generalized linear mixed effects (GLME) model to predict discrimination accuracy using linear and non-linear mixing learning parameters. We defined linear and non-linear mixing (LM and NM, respectively) as the sum and the product of orientation and color strength, respectively, on each trial. Then we multiplied these two terms by the training blocks (1, 2, …) to derive the linear and nonlinear mixed learning parameters to be used in the GLME model (see Methods for details). These models revealed significant LM and NM effects in both experiments (Experiment 1, LM×Block: *β*=0.0219, *p*<10^-8^, NM×Block: *β*=0.0569, *p*<10^-79^; Experiment 2, LM×Block: *β*=0.0277, *p*<10^-12^, NM×Block: *β*=0.0327, *p*<10^-29^). NM was nearly double as large as LM in Experiment 1, while LM and NM were close in Experiment 2.

To statistically compare the two terms between experiments, we pooled their data together and factored in experiment as a categorical moderator of our main learning predictors, NM and LM. The non-linear learning term (NM×Block) significantly differed between experiments (NM×Block×Exp: *β*=0.0244, *p*<10^-8^), while the linear learning term did not (LM×Block×Exp: *β*=-0.0061, *p*=0.2558). Importantly, this difference in NM favored Experiment 1 (reflected by positive *β* value), pointing to the more effective contribution of nonlinear term in Experiment 1 vs. 2, while LM did not differ between the two experiments (non-significant, small *β*).

Overall, these results suggest that the improved conjunction learning in Experiment 1 relative to Experiment 2 may be credited to a non-linear combination of color and orientation information which could be provided by nonlinear mixed selectivity neurons incorporated by global feedback.

### Separate feature leaning (Experiment 3)

If two features need to be updated in parallel in Experiment 2, this could hypothetically incur higher cognitive load and thus noisier credit assignment than a single, global update in Experiment 1. E.g., orientation tuned neurons could mistakenly be down-weighted if even if feedback only indicated a wrong color choice. To address this alternative hypothesis, we conducted another experiment (*n* = 12) using the same stimuli, but a slightly modified task design. Specifically, subjects saw colored gratings as in Experiment 1 and 2, but were post-cued to respond only to color or orientation on a given trial (Fig. 2e). This way, only one feature update was required on each trial, mitigating the potential difference in load between Experiment 1 and 2. Subjects showed significant learning, as evidenced by block-by-block improvements in LI (orientation: *F*(6,66)=45.54, *p*<10^-15^, *η^2^*=0.8054; color: *F*(6,66)=40.235, *p*<10^-15^, *η^2^*=0.7853). If orientation-color conjunction learning was the result of summing color and orientation channels, separate training in Experiment 3 should now yield the same results as Experiment 1 when tested on the conjunction task. We thus tested participants after training with post-cueing on the orientation-color conjunction discrimination task (identical to Experiments 1 and 2). A subset of participants (*n*=6) was briefly (180 trials) exposed to conjunction training of Experiment 1 just before this test, while the remaining subjects did not receive additional training. We found a statistically significant difference between Experiments 1 and 3 (Wilcoxon rank-sum test, *Z*=2.0892, *p*=0.0367, *d*=-0.7277), but not between Experiments 2 and 3 (*Z*=1.4608, *p*=0.1441, *d*=-0.5767) in the conjunction test (Fig. 4a). Brief additional training on the conjunction task had no statistically significant effect (Fig.4 a, *p*=1, *d*=0.1997). These results suggest that the benefit in conjunction learning that we found with global feedback in Experiment 1 relative to feature-specific feedback in Experiment 2 is not due to inefficient credit assignment in the latter experiment.

**Fig. 4.**
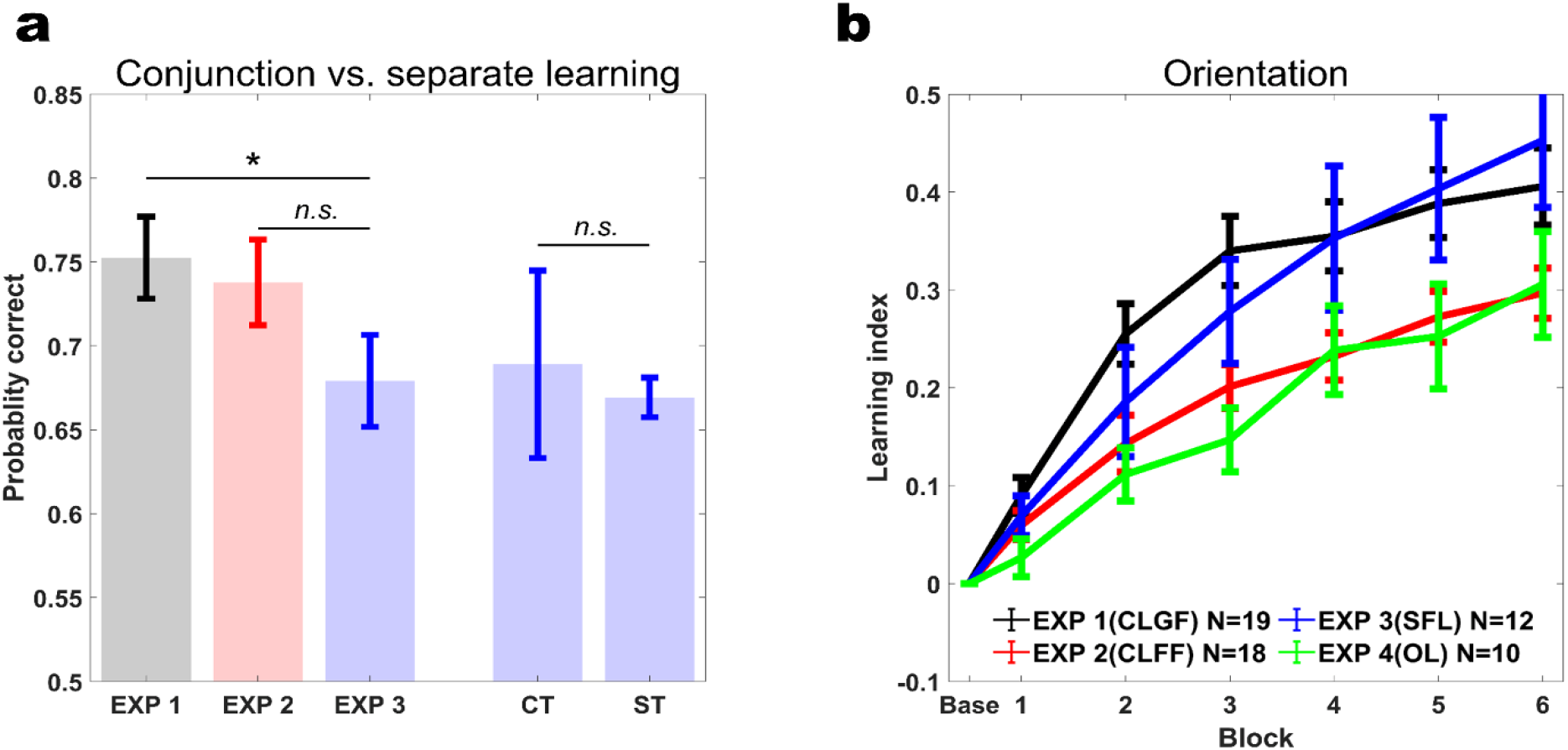
Comparing learning across experiments. **a,** Conjunction discrimination performance is significantly better after conjunction learning with global feedback (Experiment 1) than separate feature learning (Experiment 3) (*p*=0.0367, *d*=-0.7277; left bars). In contrast, there is no statistically significant difference in conjunction discrimination performance after training with feature-specific feedback (Experiment 2) versus separate feature training (*p*=0.1441, *d*=-0.5767). Brief conjunction discrimination training (identical to Experiment 1) does not significantly affect conjunction discrimination performance after separate feature learning (*p*=1, *d*=0.1997; right bars, CT: Conjunction training, ST: Separate feature training). * denotes *p*<0.05, *n.s*. denotes non-significant. **b,** Orientation LIs. Orientation is learned faster and better when trained in conjunction with color (experiment 1) than in the presence of task irrelevant color (experiment 4; *F(1,27)*=8.2049, *p*=0.008, *η^2^*=0.2331). CLGF: Conjunction Learning with Global Feedback, CLFF: Conjunction Learning with Feature-specific Feedback, SFL: Separate Feature Learning, OL: Orientation Learning. Error bars represent the standard error of the mean.

We also compared the degree of nonlinear mixing between experiments. We built a GLME model similar to the ones we had used to establish NM effects in Experiment 1, but this time constrained the model to include only the NM term and the experiment factor predicting conjunction discrimination accuracy in the test session. We find a significantly larger NM coefficient in Experiment 1 compared to the other two experiments (NM×Exp2: *β*=-0.072, *p*=0.0001, NM×Exp3: *β*=-0.08, *p*<10^-4^). Hence, nonlinear mixing of features can account for the improved conjunction learning in Experiment 1 with global feedback, relative to feature-specific feedback in Experiments 2 and 3, even for conjunctions of features thought to be as separated as orientation and color.

### Single feature learning (Experiment 4)

So far, we explored orientation learning in a context where color was task-relevant and therefore equally attended. Moreover, the learning differences among experiments we presented so far were more strongly expressed in orientation than in color, the seemingly more challenging feature to learn. We now sought to establish a baseline for single feature orientation learning while color was task-irrelevant and should thus be ignored (*n*=10). Similar to the other experiments, we drew six color levels from red to purple spectrum; however, they were not derived from individually determined psychometric functions but were set to the values with the largest distance from each other in the color space. This was done to render color as salient, and therefore distracting, as possible. We find that orientation learning in the presence of task irrelevant color was slower in Experiment 4 than in Experiments 1 (Experiment: *F*(1,27)=8.2049, *p*=0.008, *η^2^*=0.2331, Block: *F*(6,162)=123.5437, *p*<10^-15^, *η^2^*=0.8353, Experiment×Block: *F*(6,162)=6.2979, *p*<10^-5^, *η^2^*=0.1891) and Experiment 3 (Experiment: *F*(1,20)=3.6287, *p*=0.0713, *η^2^*=0.1536, Block: *F*(6,120)=63.6286, *p*<10^-15^, *η^2^*=0.761, Experiment×Block: *F*(6,120)=4.2144, *p*=0.0007, *η^2^*=0.174), but not Experiment 2 (Experiment: *F*(1,26)=0.4251, *p*=0.5201, *η^2^*=0.0161, Block: *F*(6,156)=90.1576, *p*<10^-15^, *η^2^*=0.7928, Experiment×Block: *F*(6,156)=1.0451, *p*=0.3983, *η^2^*=0.0386; Fig. 4b). This suggests that orientation learning is highly efficient when learned in conjunction with color under global feedback conditions (Experiment 1), and when attentional resources are explicitly devoted to orientation (Experiment 3), but not when highly salient color varies randomly from trial to trial and acts as a distractor (24).

### Location transfer

So far, we have appraised various learning strategies for conjunction learning and elaborated on possible neural resources recruited to serve this purpose. Now, we take another approach to specify the neural resources conducive to conjunction learning by investigating the underlying spatial receptive field (RF) properties. To this end, we test all subjects at new spatial locations after their initial training in a location transfer session. If the RFs of the trained neurons overlap with the new location, we should observe full transfer of the learning effects to this new location. In contrast, if RFs are limited in size and do not overlap with the new location, we should see full or at least partial loss of the training effects when the location is changed. Building further on this logic, we can dissociate whether conjunction learning is carried by a single population of mixed selectivity neurons, or whether pure orientation and color tuned neurons, respectively, are (also) involved. This is because color coding neurons have larger RFs than orientation coding neurons in early visual cortex of monkeys (5,25) and humans (26–28). Hence, equal spatial transfer results for orientation and color would suggest an involvement of a single population of mixed selectivity neurons (with equally sized RFs for color and orientation), while differential transfer for color and orientation would suggest an involvement of color and orientation tuned neurons (with different RF sizes for the two features). To capture potential differences in RF size, we tested transfer to a near (3 dva) and a far (6 dva) transfer location (at iso-eccentricity, Fig. 5a) in Experiment 1 and 2; in Experiments 3 and 4, we only tested the far transfer condition. We quantified transfer using a Transfer Index (TI), comparing performances between transfer and test sessions (see Methods for details). A TI of 0 indicates no transfer of learning, while a TI of 1 indicates full transfer to the new location.

**Fig. 5.**
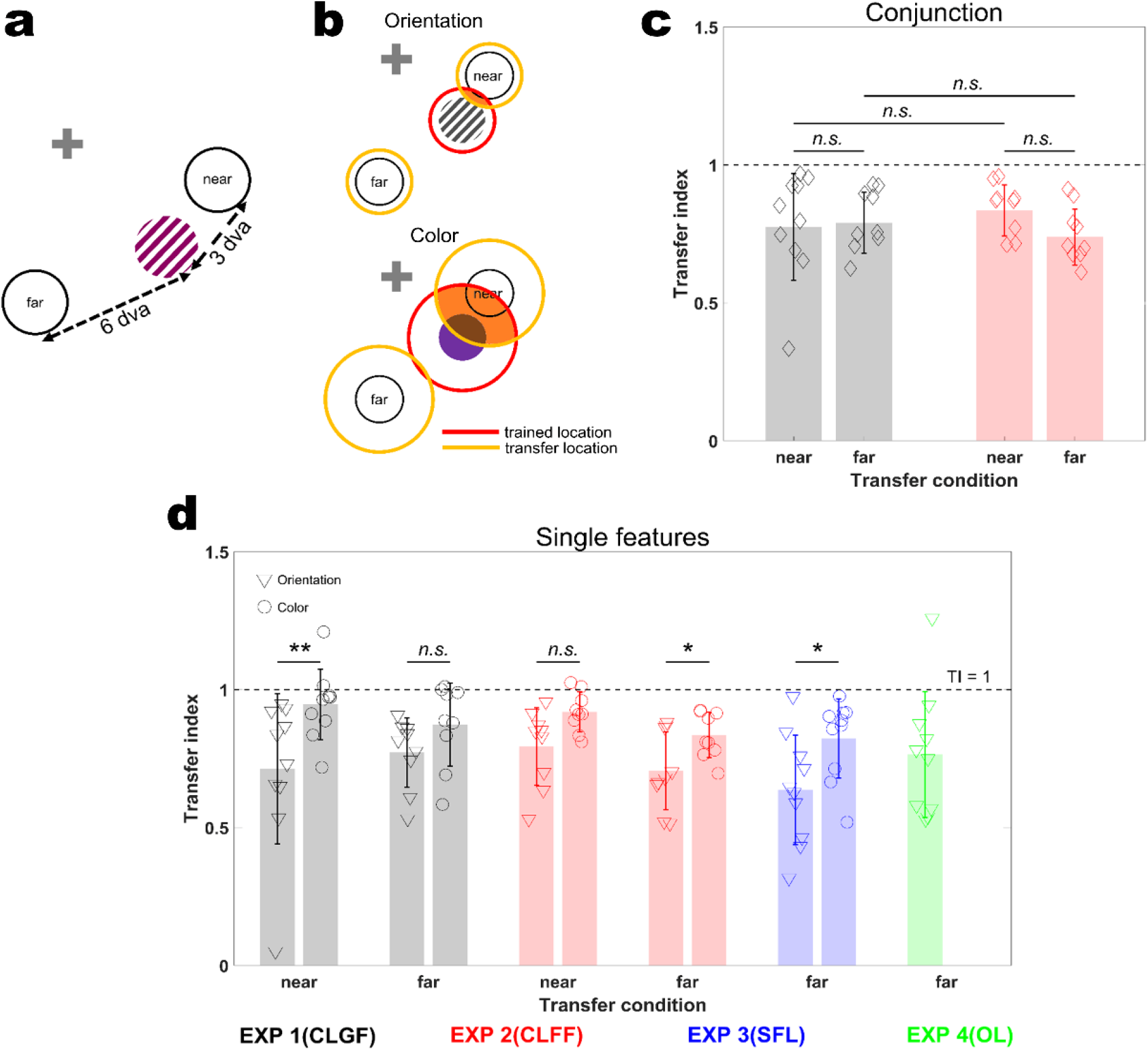
Location transfer results. **a,** Schematic illustration of the two transfer locations, near and far (3 and 6 dva, respectively, from the training location). **b,** Differential receptive field (RF) sizes of pure orientation and color selectivity neurons would lead to different transfer results for the two features: larger color RFs entail larger overlap between trained and transfer locations (area marked in orange), and hence larger TIs compared to orientation. In contrast, mixed selectivity neurons have identical RFs for color and orientation. This predicts no difference in Tis between the two features. Red circles: trained location RFs, yellow circles: transfer locations RFs. **c,** Transfer Indices (TIs) for conjunction discrimination. There was no significant difference between locations (near and far) and experiments (Location: *F*(1,33)=1.6076, *p*=0.2137, *η^2^*=0.0465; Experiment: *F*(1,33)=0.1302, *p*=0.7206, *η^2^*=0.0039; Location×Experiment: *F*(1,33)=1.2071, *p*=0.2798, *η^2^*=0.0353). **d,** TIs for single features. Orientation and color TIs are significantly different from each other (Experiment 1, near TIs: *p*=0.002, *d*=1.1, far TIs: *p*=0.1641, *d*=0.73; Experiment 2, near TIs: *p*=0.0547, *d*=1.13, far TIs: *p*=0.0273, *d*=1.1232, Experiment 3, *p*=0.0195, *d*=1.0731). Dashed black lines in panels b and c show full transfer (TI=1). CLGF: Conjunction Learning with Global Feedback, CLFF: Conjunction Learning with Feature-specific Feedback, SFL: Separate Feature Learning, OL: Orientation Learning. Error bars represent standard deviation. ** denotes *p*<0.01, * denotes *p*<0.05, *n.s*. denotes non-significant.

For conjunctions, we found partial transfer (Experiment 1, near TI: *n*=10, mean=0.7744, SD= 0.1935,*p* vs 0 <10^-6^, far TI: *n*=9, mean=0.7903, SD=0.1102,*p* vs. 0 <10^-7^; Experiment 2, near TI: *n*=9, mean=0.8347, SD=0.0921,*p* vs. 0 <10^-8^; far TI: *n*=9, mean=0.7385, SD=0.1016,*p* vs. 0 <10^-7^). TI did not significantly vary across transfer locations (near vs. far) and experiments (two-way ANOVA; Location: *F*(1,33)=1.6076, *p*=0.2137, *η^2^*=0.0465, Experiment: *F*(1,33)=0.1302, *p*=0.7206, *η^2^*=0.0039, Location×Experiment: *F*(1,33)=1.2071, *p*=0.2798, *η^2^*=0.0353; Fig. 5c). Orientation TIs followed a similar pattern (Fig. 5d), with partial transfer across locations but no difference between them (Experiment 1, near TI: *n*=10, mean=0.7129, SD= 0.2717,*p* vs. 0 <10^-4^, far TI: *n*=9, mean=0.7722, SD=0.1256,*p* vs. 0 <10^-7^; Experiment2, near TI: *n*=9, mean=0.7932, SD=0.1407,*p* vs. 0 <10^-6^, far TI: *n*=9, mean=0.7057, SD=0.1407, *p* vs. 0 <10^-7^). Moreover, there was no systematic difference in TIs for orientation between Experiment 1 and 2 across locations (Location: *F*(1,33)=0.4298, *p*=0.5166, *η^2^*=0.0128, Experiment: *F*(1,33)=0.0904, *p*=0.7656, *η^2^*=0.0027, Location×Experiment: *F*(1,33)=0.4959, *p*=0.4862, *η^2^* =0.0148), and across all four experiments at the far transfer location (one-way ANOVA; *F*(3,34)=0.9548, *p*=0.4252, *η^2^*=0.0777). For color, however, a different pattern emerged. Here, we found practically full transfer in the near (Experiment1, mean=0.9462, SD=0.1273,*p* vs. 1=0.2145; Experiment 2, mean=0.9195, SD=0.0721, *p* vs. 1=0.01), but not the far location (Experiment 1: mean=0.8731, SD=0.1499, *p* vs. 1=0.0347; Experiment 2: mean=0.8351, SD=0.0822, *p* vs. 1=0.0003). Consequently, color and orientation TIs were significantly different especially at the near transfer location, with color having larger TIs than orientation (two-sided sign-rank test, Experiment 1, near TIs: *n*=10, *p*=0.002, d=1.1, far TIs: *n=9, p*=0.1641, *d*=0.73; Experiment 2, near TIs: *n=9, p*=0.0547, *d*=1.13, far TIs: *n=9, p*=0.0273, *d*=1.1232; Fig. 5d). The same held true for Experiment 3 (*n*=10, *p*=0.0195, *d*=1.0731). This can be attributed to larger RFs for color than orientation (Fig. 5b). These results suggest that pure selectivity neurons (with different RFs for color and orientation, respectively) also partake in conjunction learning. Together with our earlier results on non-linear mixing in conjunction learning, this is most in line with a global pooling strategy in which pure color, pure orientation, and mixed selectivity neurons are combined to give rise to choice information.

### Learning effects on constituent features (pre-post training comparison)

In the preceding sections, we have shown that global feedback (Experiment 1) favors nonlinear integration of color and orientation during conjunction learning, speaking to an involvement of non-linearly mixed selectivity neurons in this learning process. Moreover, we have shown on the basis of differential location transfer results for color and orientation that purely color and orientation coding neurons also contribute to conjunction learning. We now sought to compare learning effects on pure selectivity neurons among all experiments (1 to 4). To this end, we repeated the orientation and color discrimination task we used to determine perceptual difficulty levels on the first day (pre) after completion of test and transfer tasks on the final day (post). Unlike in the main experiments, in which stimuli were defined by both features, orientation and color, only one feature was present in each task for the pre and post measurements. Therefore, performance on the pre/post tasks is diagnostic in determining the plasticity of pure selectivity neurons.

We found statistically significant learning effects across all experimental groups for orientation (two-sided sign-rank test, Experiment 1, *n*=19, *z*=3.7429, *p*=0.0002, *d*=1.9544; Experiment 2, *n*=18, *z*=3.071, *p*=0.0021, *d*=1.119; Experiment 3, *n*=10, *p*=0.002, *d*=2.0685; Experiment 4, *n*=10, *p*=0.002, *d*=2.3461; Fig. 6a). Similarly, we find improvements for color in Experiments 1, 2 and 3 (Experiment 1, *z*=2.2337, *p*=0.0255, *d*=0.7; Experiment 2, *z*=3.1953, *p*=0.0014, *d*=1.2995; Experiment 3, *p*=0.0195, *d*=1.2632) but not Experiment 4 (*p*=0.4258, d=0.4684; Fig. 6b), where color was irrelevant and should be suppressed (29). These improvements did not differ significantly across experiments (mixed repeated measure ANOVA; Orientation, Timepoint: *F*(1,53)=132.1539, *p*<10^-15^, *η^2^*=0.7286, Experiment: *F*(3,53)=0.2227, *p*=0.8802, *η^2^*=0.0124, Timepoint×Experiment: *F*(3,53)=1.7039, *p*=0.1774, *η^2^*=0.088, Color, Timepoint: *F*(1,53)=24.9849, *p*<10^-5^, *η^2^*=0.3417, Experiment: *F*(3,53)=0.9902, *p*=0.4046, *η^2^*=0.0531, Timepoint×Experiment: *F*(3,53)=1.3723, *p*=0.2613, *η^2^*=0.0721).

**Fig. 6.**
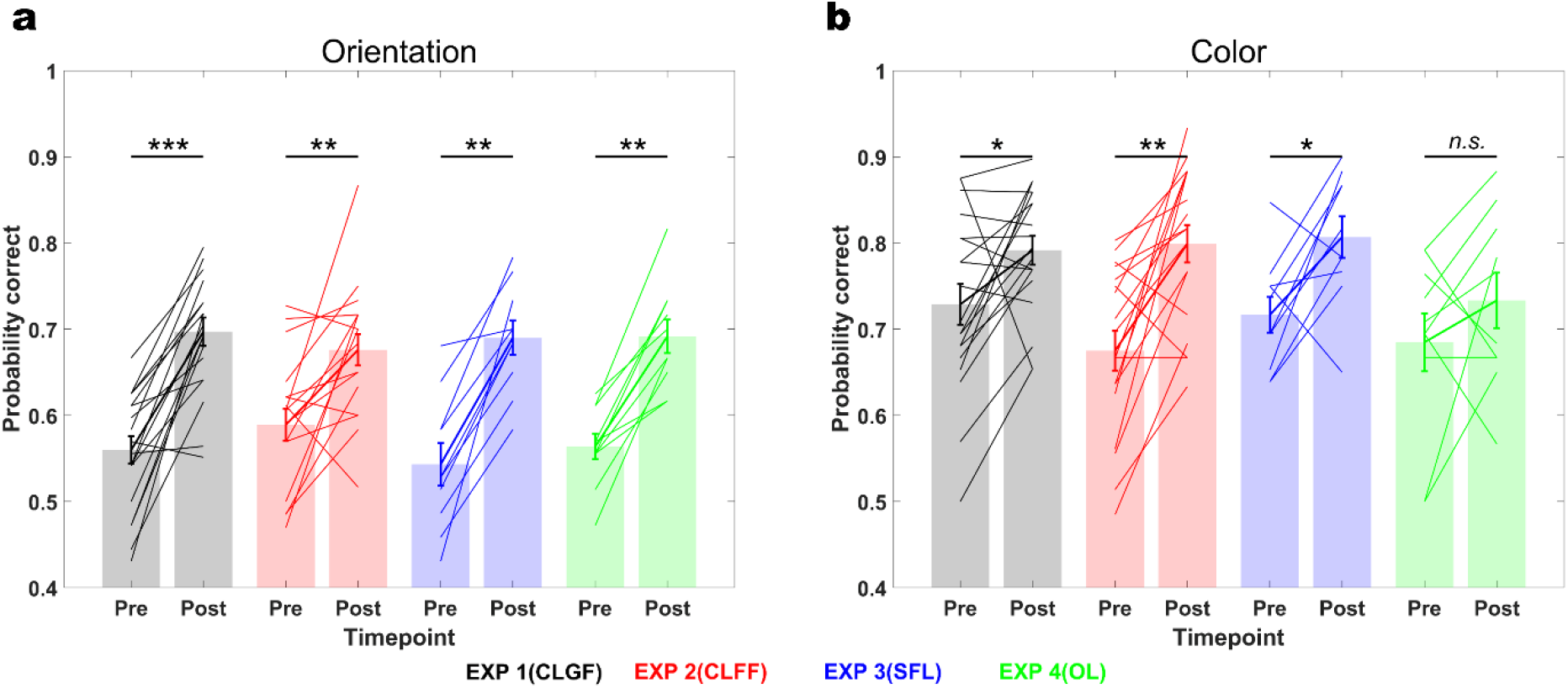
Learning effects on constituent features, pre-post comparison. **a,** Orientation discrimination performance significantly improved after training in all experiments (Experiment 1, *p*=0.0002, *d*=1.9544; Experiment 2, *p*=0.0021, *d*=1.119; Experiment 3, *p*=0.002, *d*=2.0685; Experiment 4, *p*=0.002, *d*=2.3461). **b,** Likewise, color discrimination performance improved in all experiments where color was task-relevant (Experiment 1, *p*=0.0255, *d*=0.7; Experiment 2, *p*=0.0014, *d*=1.2995; Experiment 3, *p*=0.0195, *d*=1.2632), but not in Experiment 4 where only orientation was trained (*p*=0.4258, *d*=0.4684). CLGF: Conjunction Learning with Global Feedback, CLFF: Conjunction Learning with Feature-specific Feedback, SFL: Separate Feature Learning, OL: Orientation Learning. Error bars represent the standard error of the mean. *** denotes *p*<0.001, ** denotes *p*<0.01, * denotes *p*<0.05, *n.s*. denotes non-significant.

This result provides another piece of evidence that pure selectivity neurons were also involved in training in all experiments. However, their involvement did not differ between experimental manipulations. This implies that significantly better learning in Experiment 1 compared to the others is due to non-linear mixing of color and orientation, and not learning effects on color and orientation by themselves.

### Transfer to XOR task

The finding that non-linear mixed selectivity plays an important role in conjunction learning is largely based on GLME models showing that non-linear mixing differs significantly between Experiments 1 and 2. We attributed this more efficient updating of mixed selectivity neurons in Experiment 1 than in Experiment 2 to our feedback manipulation. If non-linear mixed selectivity neurons emerge as a result of training with global feedback in Experiment 1 but not in Experiment 2, this leads to another prediction: namely, that subjects trained with global feedback should be at an advantage in solving a non-linearly separable task based on color and orientation compared to subjects trained with feature-specific feedback, because they already have representations that could support task performance in this case.

We tested this prediction by assessing how well subjects previously trained with global or feature-specific feedback in Experiments 1 and 2, respectively, could transfer learning effects to an exclusive or (XOR) task. In this 2AFC task, subjects had to respond to red-clockwise or purple-anticlockwise with a response to the right target and purple-clockwise or red-anticlockwise with a response to the left target (Fig. 7a). As in the original conjunction learning task that we used in Experiments 1 and 2, our XOR task required detection of conjunctions but, unlike in the original, linearly separable task, it required non-linear mixing of color and orientation for successful task performance.

**Fig. 7.**
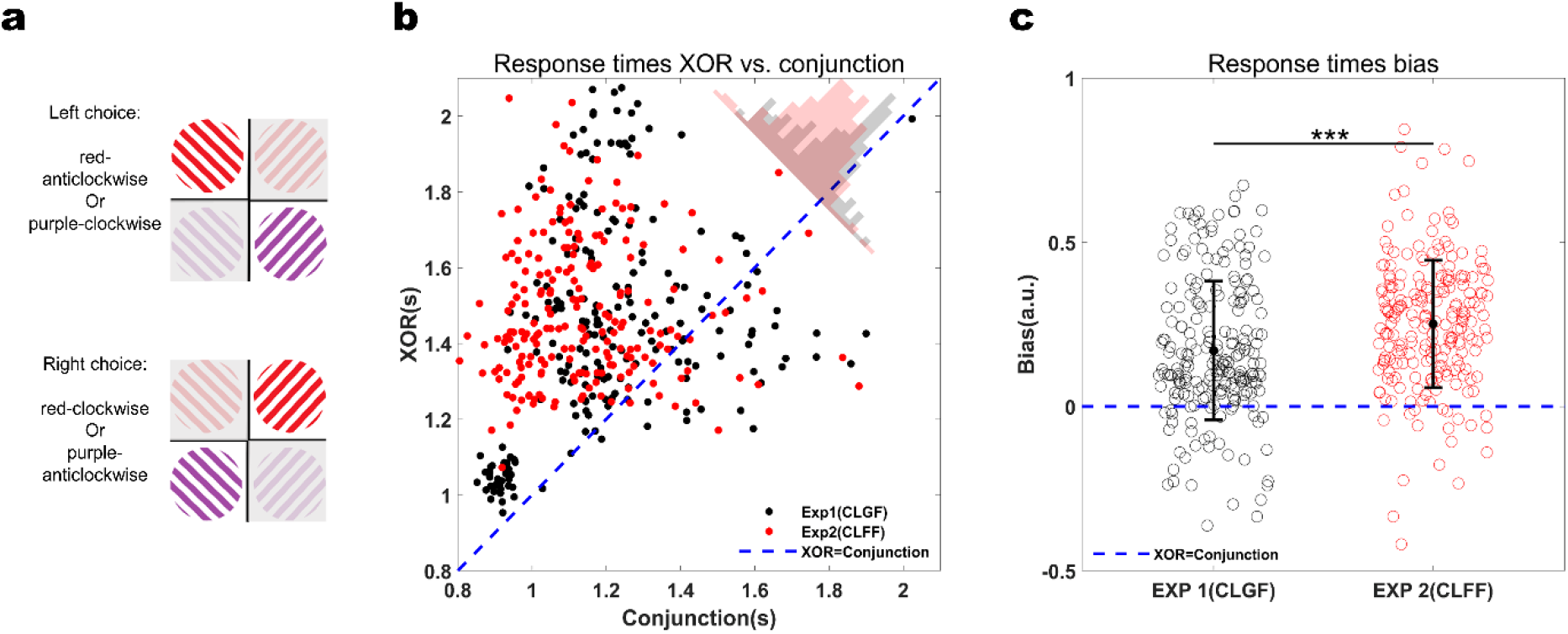
Response time (RT) comparison between XOR and conjunction tasks. **a,** In the XOR transfer task, subjects had to respond with an eye movement to the left for red-anticlockwise and purple-clockwise stimuli, and to the right for red-clockwise and purple-anticlockwise stimuli. Colors and orientations in the XOR task were identical to the ones used in the conjunction task (and are only displayed for illustration purposes here). **b,** Response times (RT) in the XOR task were generally slower than in the conjunction task (RT bias toward XOR), but this effect was larger in Experiment 2 than in Experiment 1 (mixed repeated measure ANOVA; Experiment: *F(1,10)*=0.0072, *p*=0.934, *η^2^*=0.0007, Task: *F(1,850)*=763.4, *p*<10^-15^, *η^2^*=0.4732, Experiment×Task: *F(1,850)*=26.72, p<10^-6^, *η^2^*=0.0305). Dashed blue line signifies RT equality (XOR=conjunction). Histograms of RT biases (distance from RT equality XOR=conjunction) are shown in the top right corner (black: Experiment 1, red: Experiment 2). **c,** Biases from the equality line where RT in XOR=RT in conjunction (bias=0, signified by dashed blue line). The bias towards XOR is significantly larger in Experiment 2 than in Experiment 1 (*p*<10^-5^). Error bars represent standard deviation. *** denotes *p*<0.001.

Six subjects from Experiment 1 and 2, respectively, we re-invited to perform the XOR task approximately nine months after they had finalized training. Both groups had retained more than 80% of the performance level they had attained on the original conjunction task (Timepoint TIs (Conjunction test vs. Reminder); Exp1: mean=0.8716, SD=0.0641, Exp2: mean=0.818, SD=0.1889, Experiment 1 vs. 2 rank-sum test: *p*=0.3939, *d*=-0.38). We then analyzed differences in accuracy between the original conjunction test and the XOR task using TIs between tasks (which accounts for the differences in chance levels of the two tasks). We found that both groups performed almost equally well on the XOR task in terms of accuracy (Task TIs (Conjunction test vs. XOR), Exp1: mean=0.55, SD=0.0647, Exp2: mean=0.4623, SD=0.1258, Experiment 1 vs. Experiment 2 rank-sum test: *p*=0.132, *d*=-0.8753). However, the degree of non-linear mixing (NM, as assessed using a GLME model akin to the one used to compare Experiments 1 and 3, see Methods) was significantly larger for the group trained with global feedback than the group trained with feature-specific feedback (Experiment 1 as a reference; NM: *β*=0.2332, *p*<10^-23^, Experiment: *β*=-0.24, *p*=0.2033; NM×Experiment: *β=-* 0.1035, *p*=0.0003). We then considered reaction times (RTs) differences on the Conjunction and XOR task. This is because it is well known that accuracy by itself does not necessarily differentiate performance in linearly from performance in non-linearly separable tasks (30). However, RTs can provide additional insight whether subjects perform non-linear tasks by solving them for two features sequentially. We compared RTs between groups for each color-orientation difficulty level combination (36 RT values per subject; Fig. 7b). We found significant differences in how quickly subjects with different training histories could execute the conjunction and XOR tasks, respectively (Experiment: *F*(1,10)=0.0072, *p*=0.934, *η^2^*=0.0007, Task: *F*(1,850)=763.4, *p*<10^-15^, *η^2^*=0.4732, Experiment×Task: *F*(1,850)=26.72, *p*<10^-6^, *η^2^*=0.0305). To further probe the RT differences between the two groups, we computed a RT bias index for the full RT profiles. This bias index was defined as the distance from equality where RT for Conjunction equals RT for XOR (see Methods). This revealed that subjects trained with feature-specific feedback were slower in the XOR task (relative to the conjunction task) than subjects trained with global feedback (Fig. 7c; *z*=-4.6177, *p*<10^-5^).

Together, these results suggests that subjects originally trained with global feedback were able to use non-linear mixing of color and orientation information to solve the XOR task, whereas subjects trained with feature-specific feedback relied on a different, slower strategy not involving non-linearly mixed selectivity.

## Discussion

Conjunction learning is indisputably an important aspect of perceptual learning particularly in realistic situations in which learning one feature, among many that define objects, would not lead to success. Nevertheless, most studies to date have primarily focused on single feature learning. In the present study, we evaluated possible mechanisms for conjunction learning which differ in the populations of neurons that they rely on. We found that conjunction learning can be accounted for by an improvement of global read out processes across neurons that code for constituent features and neurons with mixed selectivity for feature combinations. Removing or reducing the contribution of the latter population by using feature-specific instead of global feedback still enabled learning conjunctions, but resulted in less efficient learning. Simple summing of color and orientation information could not account for our results, as evidenced, e.g., by the finding that learning effects on separate features did not transfer to the conjunction task. Together, these findings highlight the role of mixed selectivity in conjunction learning.

Mixed selectivity neurons explicitly provide task-relevant information in conjunction learning tasks. They have been described in a number of brain areas across species, including in visual cortex (31–39). Mixed selectivity neurons are thought to improve decoding of information in downstream areas by increasing the dimensionality of the representational space, thus enabling additional hyperplanes for linear decoders operating in population space that can aid difficult and even non-linear discrimination problems (40–42). Furthermore, mixed selectivity massively reduces the number of decoding errors relative to pure selectivity (18). These benefits may explain why learning in Experiment 1, in which global feedback favored the incorporation of mixed selectivity neurons (Fig. 1a), exceeded learning in all other experiments. Learning was however also possible when feedback did not favor the incorporation of non-linear mixed selectivity. Theoretical studies have shown that pure selectivity neurons are sufficient to encode all individual task-relevant features under certain conditions (41). Hence, mixed selectivity neurons are not strictly required to solve conjunction tasks, but may provide important benefits for learning, even on the single feature level.

In fact, all task-relevant information could in theory be read out from non-linearly mixed selectivity neurons in the absence of neurons coding purely for the constituent features (41). Hence, a readout strategy that would rely exclusively on mixed selectivity neurons to solve the conjunction task is feasible and would at the same time minimize the number of neurons or weights that need to undergo plasticity during learning. This would be in accordance with precision pooling theories that suggest that the brain only reads out the most diagnostic neurons for a task at hand (20,43,44). Yet, we did not find strong evidence that conjunction learning relied exclusively on non-linear, conjunction encoding representations. Instead, several lines of evidence suggest that pure selectivity neurons are also involved in our task. Our findings seem thus more in line with readout theories that suggest that all active neurons are read out (21–23). One reason for this may lie in the strong separation of color and orientation coding stimuli in early visual cortex. Color and orientation are long thought to be represented by distinct populations of neurons (1). More recent studies have shown that this separation is not absolute and that color-orientation conjunction coding neurons exist, albeit in very limited numbers (~10% of the population) (12). This creates a trade-off: reading out information from a larger pool that contains a majority of neurons that do not explicitly code conjunctions, or from a small pool of highly informative neurons. Selective readout of highly informative neurons assures a high signal-to-noise ratio in principle but may be prone to noise if the readout pool is small. Furthermore, this readout strategy requires switching the population from which information is read out on a trial-by-trial basis. Uniform readout of all neurons, in contrast, may not always reach the same level of precision since suboptimally tuned neurons are also read out. However, it does not require switching the readout pool when variable stimuli are dealt with, and may be less prone to noise because of the large number of neurons involved. Our results suggest that the brain favors a more global readout policy, relying on a larger pool of neurons, at least in the case of discriminating orientation-color conjunctions.

Our results also suggest that global feedback is particularly effective in driving conjunction learning. This relates to the question of credit assignment. Credit assignment determines which units are selected for plasticity as much as the degree of plasticity that these units undergo. Theoretical models often consider credit assignment and associated synaptic weight updates as a local rather than a global process: the most informative units, and not all available units, are subject to plasticity. E.g., in Attention Gated Reinforcement Learning (45), which is a biologically plausible model of learning that has been extended to multi-feature learning (46), input units which are highlighted by attention receive the largest weight updates. Yet, we know that plasticity in the brain is governed by neuromodulators which tend to take effect on a broad scale. A recent theoretical study has proposed a learning rule based on global credit assignment, in which one error vector is computed in the output layer and then globally broadcasted back to the hidden and input layers to update weights (47). This architecture is on par with other state-of-the-art learning algorithms that rely on local weight updates. Together, this suggests that learning by global credit assignment is computationally feasible and biologically plausible.

Single feature learning, in particular of orientation, also benefited from global feedback during conjunction training. However, highly effective single feature learning was not unique to the global feedback condition during conjunction training (Experiment 1), but also occurred in Experiment 3, when subjects were post-cued to attend either color or orientation. However, Experiment 4, in which only orientation was task-relevant, and which had double the number of trials than Experiment 3, showed less improvements in orientation discrimination than Experiments 1 and 3. This may be because variability in a highly salient yet unattended feature may block or at least insert noise to credit assignments and thus impair learning. This would be consistent with the finding that top-down attention can be insufficient in suppressing highly salient color distractors (48). More speculatively, the diminished learning effects in Experiment 4 might also indicate that mixed orientation-color representing neurons exist in larger numbers and/or play a more critical role than previously thought. In this case, task-irrelevant color information would be more difficult to filter out from the readout pool and may thus have had an excess effect on orientation learning.

Another interesting finding in this study are steeper learning curves in conjunction learning with global feedback (Experiment 1) compared to the other experimental conditions (Fig. 4b), which we observed during the early training phase (days 1 and 2). Can non-linear mixed selectivity also account for this effect? In principle, if decoding accuracy for a single feature is *p*, decoding accuracy of two independent features coming from pure selectivity sources decays as *p^2^*, while decoding accuracy of mixed selectivity would not decay by the same extent, resulting in higher learning rates. Previous evidence on whether mixed selectivity, compared to pure selectivity, speeds up or slows down learning, however, is somewhat mixed. A theoretical study has shown that the larger the number of mixed selectivity neurons in a network, the higher the achievable learning rates (40). Yet, it has been argued on empirical grounds that single feature learning by pure selectivity is faster than conjunction learning by mixed selectivity, because the former can entail more frequent weight updates (49). This study explored unsupervised learning with high-dimensional stimuli and with a variable reward schedule. The authors show, in this particular environment, that subjects started learning stimulus-reward associations by attributing reward probabilities to single features even if conjunctions were more informative of the reward. Subjects then slowly converged onto conjunction associations as they explored the stimulus space in the course of training. Our study differs from this previous study in several respects, in particular because we used a fixed reward schedule, supervised learning, and low-dimensional conjunctions. Future studies explicitly geared to these parameters will be needed to determine if and when mixed selectivity expedites unsupervised conjunction learning.

It is worth reemphasizing that the benefits of global feedback that we attribute to the incorporation of mixed selectivity neurons in conjunction learning occurred for a linearly separable task. This concurs with the finding that mixed selectivity exists in the brain even when it is not strictly behaviorally relevant (18). However, we also found evidence that subject trained with global feedback had an advantage over subjects trained with feature-specific feedback in transferring learning effects to a non-linear XOR task, even several months after the initial training. XOR tasks are a canonical example of non-linear seperability (42,50). The reason is that a linear decoder can immediately readout XOR information from non-linear mixed selectivity units. A recent empirical study has shown that mixed-selectivity neurons in early visual, retrosplenial, and posterior parietal cortex of mice indeed predict accuracy on discrimination tasks on a trial-by-trial basis, underlining the biological validity of this concept (32). While non-linear mixing of color and orientation information predicted subjects’ accuracy during the XOR task in the group previously trained with global feedback, we did not observe differences in accuracy on the XOR task between the two training histories on average. However, it has been argued that when non-linear tasks such as the XOR task are verbalizable, they can be decomposed into two separate feature tasks and solved sequentially (51). E.g., in our task, subjects could first determine color and then respond to orientation. In this case, XOR tasks can be solved using only pure selectivity neurons but require prolonged deliberation time. Indeed, we observed that subjects trained with global feedback were faster in executing the XOR task than subjects trained with feature-specific feedback. This suggests that the emergence of mixed-selectivity neurons in conjunction learning with global feedback enables incorporating these units into new, non-linear tasks.

In summary, using several experimental manipulations and a comprehensive analysis, bolstered by sufficiently large sample sizes, we have evaluated several scenarios that could possibly account for conjunction learning. Among those, conjunction learning on the basis of both mixed and pure selectivity neurons under a global pooling strategy most comprehensively explained our results. Future physiological studies are required to further probe the neural underpinnings of conjunction learning and to explore the mechanistic role as well as the locus of mixed selectivity neurons in this context. Furthermore, it remains to be determined how conjunction learning affects decision making processes. A recent study investigating decision making for feature conjunctions reported that features are processed in parallel but integrated serially into one decision (52). How learning affects these processes remains a question for future studies.

## Methods

### Participants

Participants aged 18 to 45 with no previous history of neurologic/psychiatric disorders with normal or corrected to normal vision were invited for the experiment. This invitation sent via an online recruitment system of the European Neuroscience Institute Göttingen (ORSEE (53)) and hence, most of them were students or university employees. A brief D15 color vision test (54) was performed to exclude subjects with impaired color vision. In total 63 participants volunteered for the experiments. Of those, 4 failed to improve throughout training and were thus excluded from the analysis, leaving 59 participants in total (39 female, 3 left-handed, mean age 26 yrs, SD 4.16 yrs). 19 participated in Experiment 1 (13 female, 1 left-handed, mean age 24.84 yrs, SD 4.25 yrs), 18 in Experiment 2 (12 female, 1 left-handed, mean age 27.7 yrs, SD 4.82 yrs), 12 in Experiment 3 (7 female, 0 left-handed, mean age 25.8 yrs, SD 3.19 yrs), and 10 in Experiment 4 (7 female, 1 left-handed, mean age 25.2 yrs, SD 3.08 yrs). No sample size estimation was performed but sample sizes were larger or at least equal to previous perceptual learning studies. All participants were paid 8 €/hour. To maintain motivation across whole sessions, they were additionally paid by 2€/hour for any improvement from their last score. Participants were fully instructed for the experiment and gave a written informed consent before participation. All procedures were in accordance with the Declaration of Helsinki and approved by the Ethics Committee of the University Medical Center Göttingen (protocol number 29/8/17).

### Stimuli and procedure

Participant in all experiments were trained on orientation and/or color discrimination tasks (detailed below). The experiments took place over 3 to 5 consecutive days with one training session per day. Thus, training sessions were separated by one night of sleep. Stimuli were presented on an LCD monitor (ViewPixx EEG, refresh rate 120 Hz, resolution 1920 × 1080 pixel, viewing distance 65 cm) in a darkened, sound-attenuating booth (Desone Modular Acoustics). Stimulus delivery and response collection were controlled using Psychtoolbox (Brainard, 1997) running in Matlab (The Mathworks, Inc.) on Windows 10. During all experiments, we continuously acquired pupil and gaze measurements using a high-speed, video-based eye tracker (SR Research Eyelink 1000+). Participants had to report their choices with a saccade aimed at defined circular target positions on the screen. For a valid response, they had to hold their fixation for 500 ms until they heard a click sound informing them that the response was accepted. The trials with shorter fixation duration or saccades during the fixation period were aborted and repeated at the end of the block.

### Experimental design

#### Pre and post training orientation/color discrimination task

These tasks were designed to identify individual difficulty levels in orientation and color discrimination. To this end, 15 color and 15 orientation levels were displayed in a form of solid disc or square-wave gratings in the Color and Orientation Discrimination tasks, respectively (CD and OD). In the CD task, color levels were iso-luminant chromaticities sampled from the red-purple spectrum of Lab space (L=25, a= (96: 54), b= (−4: −46), luminance= 15.74 cd/m^2^; black dots in Fig. 2b). The orientation levels in OD task were gray unipolar squared-wave gratings with polar angles ranging from 36-54° (45° and 45°±Δ=[0.7°, 1.5°, 2.5°, 3.5°,5°, 7°, 9°]). Other parameters were fixed between gratings (spatial frequency=2 cpd, luminance 15.85 cd/m^2^, gray intensity=0.4). The phase of the gratings randomly took one of four values (144°,168°, 192°, 216°) on each trial. All stimuli, whether colored discs or gray gratings, spanned 3° of the visual field and was presented on the southeast diagonal at 6° eccentricity (distance from both meridians=4.24°) against a black background. Each stimulus condition repeated 13 times in the pre and 10 times in the post training sessions (described below). Responses to the first presentation of stimuli in the pre training sessions were excluded from analysis due to the probable novelty bias.

Each trial started with a fixation phase (500 ms). Then, the stimulus was presented for 200 ms, immediately followed by two circular targets which appeared either on the bottom and top side of the central cross (color task, CD), or on the right and left side of the cross (orientation task, OD) as shown in Fig. 2a. Participants were instructed to respond by a saccade directed toward either of those targets with the following codes: In the CD task, top target coded reddish and bottom target coded purplish colors. Correspondingly in the OD task, right and left targets represented clockwise or anti-clockwise tilt from the diagonal axis (45°), respectively (Fig. 2c). Participants were asked to respond as accurately and quickly as possible but without a time limit. The reference grating (45°) was presented before the initiation of the OD task for a duration controlled by the participant (to proceed to the task, they had to press a button on keyboard). The trials were separated by an inter-trial-interval (ITI) of 500 ms. Participants were free to take a break whenever they want between the trials. However, short breaks (at least 30s) were inserted every 90-140 consecutive responses to avoid oculomotor fatigue.

Psychometric curves were obtained using a Weibull fit to the response probabilities of the chromaticity and orientation levels (Fig. 2d). Based on these curves, the points in stimulus space (LAB values or polar angles) that correspond to 0.2, 0.3, 0.4, 0.6, 0.7, 0.8 response probabilities were specified. This procedure resulted in 36 chromatic gratings which varied in both orientation and color dimensions (6 orientations × 6 colors = 36). These values were later used to shape individualized stimulus spaces for the main course of training.

Both OD and CD tasks were repeated on the final day after completion of training phase to provide a measure of how orientation and color discrimination was affected by each experiment.

### Experiment 1, conjunction learning with global feedback

#### Training phase

In Experiment 1, participants had to combine orientation and color to come up with a single response. A chromatic grating was presented at 6° eccentricity while subjects were fixating on the central fixation cross. Stimulus size and location were identical to the preceding OD/CD tasks. After stimulus offset, participants faced 4 disc-shaped gray targets on the diagonals of the screen (9° away from center). They had to report the orientation-color conjunction by looking at one of those, combining the assignments of the previously performed pre-training OD/CD task: reddish top, purplish bottom, clockwise right, anti-clockwise left. E.g., if subjects perceived a reddish clockwise tilted grating, they had to direct their gaze to the top right target. Each choice was followed by audiovisual feedback informing of response correctness on the conjunction level. After correct choices, the targets color changed from gray to green accompanied with a click sound, and after error choices, the target became red accompanied by an error sound (Fig. 2e).

Training continued for four days. The first day contained 576, the second and third day 1080, and the last day 216 trials. Every ~500 trials were considered one training block in analyses as an indicator of training progress in time. This way, the first day consisted of one, and the second and third day of 2 blocks each. Training on the last day served as a reminder and not considered for the analysis. Presentation order of stimuli was randomized across trials.

#### Test phase

Following a brief training session on the final day, participants were tested on their performance on the task they had been trained for. The test session was identical to the training sessions but with no choice feedback. This session consisted of 432 trials.

#### Transfer phase

The transfer phase was similar to the test phase, but we systematically changed the location of the stimuli. Stimuli were presented in two different locations on the screen (location transfer). In the near transfer condition, stimuli were moved 3° upward. In the far transfer condition, stimuli were moved 6° in the opposite direction (Fig. 5a). Eccentricity and stimulus size were identical to the test and training sessions. Not all participants performed both transfer tasks to prevent interference effects, yet, they were randomly assigned either to the near or the far transfer groups (*n*=10 in the near, and *n*=9 in the far transfer group).

### Experiment 2, conjunction learning with feature-specific feedback

Subjects performed the same task as in Experiment 1. The only difference was the format of the feedback that was provided on each trial. Instead of global feedback (conjunction level), subjects were provided with feature-specific feedback. Following saccades to the response target, the circular disc shape of the target was changed into a half-grating half-disc shape. The left half-gratings’ color, which was vertical parallel lines, carried orientation feedback and the right half-disc’s color carried color feedback (Fig. 2e). Akin to Experiment 1, and all other experiments, green represented correct and red represented error choices.

This design aimed to enable parallel feature learning. Due to the possible higher cognitive load of dual feedback and hence incremented processing time, the ITI increased from 500 ms to 1500 ms. Test and transfer phases were similar in design to Experiment 1.

### Experiment 3, separate feature training

Stimuli were identical to Experiments 1 and 2. However, instead of training conjunction discrimination, participants were trained on single feature (orientation/color) discriminations. Orientation and color trials were half-split and randomly intermixed across the training sessions, so that it was impossible to anticipate the next trial (orientation or color). This means that each trial’s task (orientation or color) was only known after the stimulus offset depending on the post-cue target positions. If the two response targets appeared on the bottom-top sides of the screen, color had to be responded to, and otherwise, if the targets appeared on the right-left sides, subjects had to report orientation (Fig. 2e). We reasoned that this would make participants attend to, and hence process, both features during stimulus presentation, but only respond to and receive feedback on one of them. This design aimed to relieve higher cognitive load of Experiment 2 due to dual feedback structure while preserving similar input to the involved neurons in the learning process. Feedback was provided for the respective post-cued feature.

On the final day, participants were split into two groups. One group continued separate feature training (*n*=6), while the other group received conjunction training as in Experiment 1 (*n*=6). This last training session entailed 216 trials in both groups.

#### Test phase

Participants underwent two types of tests, Separate Test (ST), and Conjunction Test (CT), without feedback. The paradigm in the former was similar to training sessions (separate feature training) while the paradigm in the latter changed to the format implemented in Experiment 1, conjunction discrimination. Due to the longer duration of last day that consisted of several tasks, 3 participants performed all tasks on the 4^th^ day, while the rest of the subjects preferred to return for a 5^th^ day. It should be highlighted that in the participants who opted for 5 days and were in the conjunction training group, this training was carried out on the 5^th^ instead of the 4^th^ day while on the 4^th^ day they continued with their usual paradigm (separate feature training). Only the far location transfer was tested in this experiment.

### Experiment 4, single feature training

This experiment was about orientation training in the presence of task irrelevant variable color. Hence, only orientation had to be reported, ignoring the color dimension (Fig. 2e). Compared to the stimulus preparation of the other experiments, one major change was applied here: instead of taking the 6 color levels from the individually determined difficulty levels (OD task), they were equally sampled from the whole red-purple spectrum, keeping the largest possible inter-level distances. This was done to increase color saliency maximizing its distracting effect.

#### Test phase

As in the other experiments, the tasks in the test sessions were identical to the training session but no feedback was provided. As for Experiment 3, only the far transfer condition was tested.

### Exclusive Or (XOR) transfer

6 participants from Experiment 1 (4 female, 1 left-handed, mean age 27.33 yrs, SD 5.95 yrs), and 6 from Experiment 2 (4 female, 0 left-handed, mean age 26 yrs, SD 2.83 yrs) were invited to perform a single-session XOR task. This was done several months after they had finished their training (~11 months after Experiment 1 and ~6 months after Experiment 2). Prior to the XOR task, they were reminded of what they have learned by doing ~500 trials of the original conjunction task (with feedback). The XOR task, akin to the conjunction task, requires conjunction discrimination but with a different underlying rule. Subjects had to determine if the presented stimulus was Red-Clockwise or Purple Anticlockwise by looking at the right target, or conversely, or whether it was Purple-Clockwise or Red-Anticlockwise by a saccade aimed at the left target (Fig. 7a). Hence, this task, unlike the conjunction discrimination tasks of Experiment 1 and 2, could not be done by a linear combination of the two features.

### Statistical Analysis

#### Learning Index

Learning was measured and tracked using a baseline normalized performance index (55). For this, we first defined baseline as performance on the first 108 trials of the first training day. Then, a Learning Index (LI) was defined using following equation:

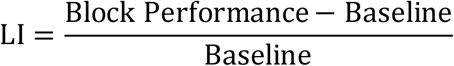

Within and between comparisons were then performed on these indices, computed separately for each block and each participant. To compare LIs between experiments, mixed repeated measure ANOVAs were performed on Aligned Ranked Transformed (ART) (56) LIs. To scrutinize difference between the two conjunction learning experiments (Experiment 1 and 2), contrast tests (57) were performed to compare baseline to each of timepoint’s LI (Blocks) across the two experiments (Baseline – Block1,2,… | Experiment 1 vs Baseline – Block1,2,… | Experiment 2).

#### GLM analysis

We used generalized linear mixed models (GLME) to determine the effect of non-linear and linear mixing on the learning curves. To define these two terms, we first defined Orientation Strength (OS) and Color Strength (CS) as a relative distance of a given feature level from chance level. Using this metric, the perceptual levels of 0.2, 0.3, 0.4, 0.6, 0.7, 0.8 yielded in relative strength values of 3, 2, 1, 1, 2, 3 respectively. Non-linear Mixing (NM) and Linear Mixing terms (LM) was dot product and sum of OS and CS values of a given stimulus in a given trial, respectively (NM=OS×CS, LM=OS+CS).

To determine learning effect on these two terms, we multiplied them by training block which was an integer from 1 to 6, giving rise to non-linear and linear learning terms (LNM=NM×Block, and, LLM=LM×Block). These predictors were then fed to a GLME model to predict conjunction response probability of each trial:

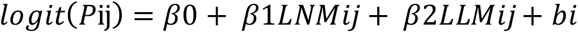

Where Pij was probability being correct (0 or 1) of subject i in trial j, *LNMij* and *LLMij* were non-linear and linear learning terms of subject i in trial j, and *bi* was the random-effects intercept for participant i that accounts for subject-specific variation.

To study the effect of experiment on either of the terms, it was factored in the model as following:

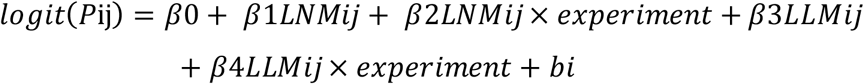

*experiment* was categorical moderator in the model. Experiment 1 served as reference. Hence, a negative coefficient means declining from Experiment 1 to 2.

To compare non-linear mixing between the conjunction test sessions of Experiment 1 to 3, we used following model:

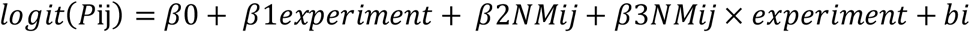

Again, Experiment 1 served as reference and *experiment* labels 1 to 3 were categorical moderators.

Finally, non-linear mixing on the XOR tasks for subjects trained in Experiment 1 and 2 were compared using following model:

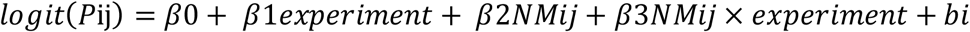

Here *Pij* was probability correct in the XOR task of participant i, trial j. Experiment 1 and 2 were the two categorical moderators and Experiment 1 was the reference for comparison.

#### Transfer Index

As in (58), the Transfer Index was defined using the following equation:

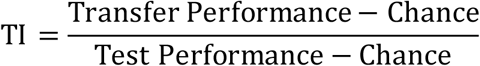

We computed TI for three transfer situations: to compare performances in the trained and untrained locations (location TI), to measure the amount of learning subjects retained after a long interval before execution of the XOR task (timepoint TI), and finally to assess the performance differences between conjunction and XOR tasks (task TI). Chance levels were 0.25 in the case of conjunction task (4AFC) and 0.5 in the other cases (XOR, single features, etc.).

#### Response Time Bias

To compare the degree of change in RT in the XOR task relative to the main Conjunction task, we defined an RT bias index in Conjunction-XOR RT space. The x-axis in this space was RT in the conjunction test, and correspondingly, the y-axis was RT in XOR task (Fig. 7b). Each point in this space represents RT to each stimulus variation in each subject in the conjunction test (x coordinate) vs. the XOR task (y coordinate). Consequently, bias index for each point in this space was defined as its distance from the diagonal which is the equity line (conjunction RT=XOR RT, Fig. 7c). Negative values signify slower conjunction response than XOR, and conversely, positive values indicate slower response to XOR.

#### Data Analysis

Unless otherwise mentioned, statistical comparisons were performed using Wilcoxon rank-sum tests to compare between two independent groups and signed-rank tests for one sample comparisons or the difference between paired samples. Before fitting ANOVAs, data were aligned and rank-transformed (56,59) using the ARTool package (version 0.11.1) (60) in R (version 3.6.1, R Core Team, https://www.R-project.org) to satisfy distributional assumptions. Contrast tests with ART were performed following the method described in (57). Reported effect sizes are Cohen’s *d* or partial *η^2^*. *Z*-statistics are provided for large samples or approximations for computing *p*-values in non-parametric tests. All analyses and model fitting were done using custom code written in MATLAB (version R2019B, MathWorks, Inc., Natick, MA) and R. The LAB color wheel displayed in Fig. 2b was produced using the Computational Color Science Toolbox in MATLAB (61).

## Data availability

The data will become publicly available upon acceptance of the paper.

## Acknowledgements

We would like to thank S. Fusi for helpful discussions. This project has received funding from the European Research Council (ERC) under the European Union’s Horizon 2020 research and innovation programme (Grant agreement No. 802482, to CMS). CMS is supported by the German Research Foundation Emmy Noether Program (SCHW1683/2-1). The funders had no role in study design, data collection and interpretation, decision to publish, or preparation of the manuscript. The authors declare no competing financial interests.

## CRediT author statement

B.K., Conceptualization, Methodology, Investigation, Formal Analysis, Visualization, Data curation, Writing – original draft preparation; C.M.S., Conceptualization, Methodology, Writing – original draft preparation, Supervision, Project administration, Funding acquisition.

